# When crayfish make news, headlines are correct but still misleading

**DOI:** 10.1101/2024.09.07.611802

**Authors:** Zen Faulkes

**Author notes:** Fredericksburg, VA, 22408, USA.

## Abstract

Crayfish are well known to many but not often newsworthy, so cases where crayfish are covered in international news provide an example of how science journalism covers a news story about basic research. Headlines have a disproportionately large influence on people’s factual knowledge and perceptions of stories covered in media. I tracked online media coverage of one scientific paper involving marbled crayfish and analyzed the headlines used by the articles. Articles were framed as “news,” but almost no headlines contained “new” facts that first appeared in the target scientific paper. The fact that appeared in the most headlines (that marbled crayfish reproduce by cloning) was over a decade old. Headlines misled readers into thinking a “breakthrough” was made by one team, rather than showing incremental advances by many teams of researchers over years.

## INTRODUCTION

Headlines have the potential to be disproportionately influential in shaping people’s knowledge and perceptions. Headlines are designed to grab attention, but many people will only read the headline and not the article that follows (the replacement effect) (Tannenbaum 1953; Condit et al. 2001). The function of headlines is to grab attention, so there is the possibility that the content of the headline differ from the rest of the article (Andrew 2007), and that will influence the perception of the main article (the framing effect) (Pfau 1995; Condit et al. 2001; Ecker et al. 2014).

There is significant concern about how reporting of science conveys matters related to risk (Cassidy 2006). For example, flawed reporting of medical research can lead people to make poor decisions, which in turn impacts both personal and public health. But there are many other branches of science where the stakes are different. The direct relevance to people’s lives may be less clear than personal health or wealth, but they receive media attention and are considered newsworthy because they are new and interesting.

Crayfish can provide an example of a “low stakes” science story. Crayfish are familiar to many people, either as food, local wildlife, or aquarium pets, but unlikely to recognize astacology as a scientific discipline. Since 2007, I have curated the website Marmorkrebs.org (http://marmorkrebs.org). This site is specifically devoted to the parthenogenetic marbled crayfish, Marmorkrebs (*P. fallax* f. *virginalis* (Martin et al. 2010a) or *Procambarus virginalis* (Lyko 2017)), which is both an emerging model organism (Vogt 2008; 2011; Faulkes 2016) and unwanted non-indigenous crayfish (Jones et al. 2009; Kawai et al. 2009; Nonnis Marzano et al. 2009; Chucholl and Pfeiffer 2010; Martin et al. 2010b; Chucholl et al. 2012; Bohman et al. 2013; Samardžić et al. 2014; Faulkes 2016; Pârvulescu et al. 2017). This Marmorkrebs.org website contains a bibliography of all scientific articles I am aware of that make substantial references to marbled crayfish. I keep watch of news concerning this species for material for a bog associated with the website, and I previously tracked online references to this species, and found most mentions were from pet owners selling marbled crayfish (Faulkes 2013). Thus, I had both a clear timeline of how research on this species has developed, and experience with baseline levels of online discussion. But a new paper published in 2018 (Gutekunst et al. 2018) received significantly more media attention than any previous paper about this crayfish species. At the time the paper was released, the Altmetric scores for other papers about marbled crayfish in my bibliography was typically less than ten, with one paper barely having a score above a hundred. But within weeks, the Altmetric score for (Gutekunst et al. 2018) paper rose to over 700. This research paper, unusually well covered for the discipline of astacology, provided an opportunity to see how a basic science story is covered in media. I wanted to assess the accuracy and emphasis of news headlines for articles that were prompted by a scientific paper about marbled crayfish (Gutekunst et al. 2018).

## METHODS

The focal paper was Gutekunst et al. (2018). This paper was embargoed before release, which allowed some journalists to get advance copies of the article, press release, and an opportunity to write stories that were timed to coincide with the release of the paper. I had already created alerts for “marbled crayfish” and “Marmorkrebs” on Google Alerts (https://www.google.com/alerts), which I used for creating content on the Marmorkrebs.org website. After the paper was published, I also checked the article’s Altmetric page (https://www.altmetric.com/details/32694958) for links to news coverage. I then compiled a list of online articles covering or responding to the Gutekunst et al. (2018) paper, including the initial press release and a post by the senior author on the journal’s blog. The list included the date of publication, headline in English, author (if given), the website name, and URL. If a headline was given in another language, it was translated to English using Google Translate (https://translate.google.com). This compiled data file was deposited on Figshare (https://doi.org/10.6084/m9.figshare.6211082). Articles were archived as PDF files using Web2PDF (http://web2pdfconvert.com) or an HTML file.

## RESULTS

Headlines largely ignored the two major new results in Gutekunst et al. (2018) (Table 1). The first major result was the publication of the first complete decapod crustacean genome. No headlines mentioned the genome sequence or words such as “gene” or “genetic” (0%; 0 of 134). The second major result was documenting the rapid spread of Marmorkrebs in Madagascar since they were found there in 2007 (Jones et al. 2009; Kawai et al. 2009). Only two headlines (1.5%, 2 of 134) mentioned the spread of marbled crayfish in Madagascar, if one includes a headline that referred to “Africa.” More headlines mentioned the spread of marbled crayfish to Europe (12.7%, 17 out of 134) than Madagascar. The spread of marbled crayfish in Europe is well documented (Chucholl and Pfeiffer 2010; Martin et al. 2010b; Chucholl et al. 2012), but no data about this was presented in (Gutekunst et al. 2018). Even more headlines stated that marbled crayfish were spread “the world” (18.7%; 25 of 134). While it is true that marbled crayfish have been found in the wild on three continents, their presence in Africa is limited to Madagascar(Jones et al. 2009; Kawai et al. 2009), and their presence in Asia is limited to a few documented introductions in Japan (Kawai and Takahata 2010; Fujiie 2017). Taking these together, 32.8% (44 of 133) headlines made mention of the marbled crayfish as an introduced and potentially invasive species.

**Table 1.** Timeline of marbled crayfish discoveries and mentions in 2018 headlines.

| Marbled crayfish fact | Year demonstrated | Mentions in 2018 headlines |
| --- | --- | --- |
| Asexual reproduction | 2003 (Scholtz et al. 2003) | 14.2% |
| Clonal reproduction | 2007 (Martin et al. 2007) | 57.5% |
| Introduced into Europe | 2010 (Chucholl and Pfeiffer 2010; Martin et al. 2010b) | 12.7% |
| Probable mutant origin | 2015 (Vogt et al. 2015; Martin et al. 2016) | 39.6% |
| First complete genome | 2018 (Gutekunst et al. 2018) | 0.0% |
| Spread through Madagascar | 2018 (Gutekunst et al. 2018) | 1.5% |

The press release headline focused on the prospect of using marbled crayfish as a model for the study of cancer. But few headlines followed the example of the press release. Only 6.0% (8 of 134) of headlines included the words “cancer” or “tumor,” including the original press release.

The most common piece of information in headlines (61.9%; 83 of 134) was some reference to the reproduction in this species. This included 57.5% (78 of 134) headlines that included some variation of the word “clone.”References to asexual reproduction was less common, and often appeared in conjunction with some variation of the word “clone” (12 included “asexual” and “clone”). Three included the word “asexual” but not “clone”; two described the species as “all-female” but not as “clones” or “asexual; and one used the word “parthenogenetic.” There was substantial overlap in terms used related to reproduction, with words like “all-female” and “clones” appearing in the same headline. That marbled crayfish are all female and reproduce asexually was shown in the first English language paper about marbled crayfish in 2003 (Scholtz et al. 2003). “Asexual” is not synonymous with “clone,” however (Martin 2016). That marbled crayfish truly clone themselves; i.e., have offspring that are the same as the mother genetically; was demonstrated later (Martin et al. 2007).

Most headlines gave an indication of the type of animal being reported on. “Crayfish” or “crawfish” appeared in 85.1% of articles (114 of 134), and 1.5% (2 of 134) used a more generic term for crustacean. Of the remaining 13.4% (18 of 134) gave no indication as to the type of animal discussed by the news, but half of those (9 of 18) included marbled crayfish as part of compilations or round-ups of news stories. The remainder used generic words like “clone,” “species,” or “creature.”

Words like “mutant,” “mutation,” and “mutated” appeared in 39.6% of headlines (53 of 134). The origin of marbled crayfish was unknown when it was discovered, because there was no wild population. The two leading hypotheses for its origin was hybridization and mutation. Two labs concluded independently that marbled crayfish likely arose by genome duplication (allopolyploidy) (Vogt et al. 2015; Martin et al. 2016), which is consistent with genetic data presented in Gutekunst et al. (2018).

A few headlines (5.2%; 7 out of 134) referenced that marbled crayfish were first noticed as in the pet trade, by using the words “pet” or “aquarium.”

As noted above, some headlines used artistic license or exaggeration, like describing marbled crayfish as “taking over world.” But only one headline was factually wrong. It read, “Mutant fish that CLONES itself and is taking over rivers worldwide leaves experts baffled.”

First, crayfish are not fish. Second, Gutekunst et al. (2018) showed that experts were informed, not “baffled.” (One other article described scientists “scratching their heads.”) There may have been reason to describe biologists as “baffled” in 2003 when marbled crayfish were first introduced to the scientific literature (Scholtz et al. 2003). But there is little reason to call biologists “baffled” in 2018, when the major questions about how and why marbled crayfish reproduce asexually were mostly answered by the intervening fifteen years of research.

The scientific article and accompanying press release was released Monday, 5 February 2018. Most articles (73.9%; 99 of 134) appeared within the following week (Figure 1). But there was a long tail. One story appeared 86 days after the release date.

**Figure 1.**
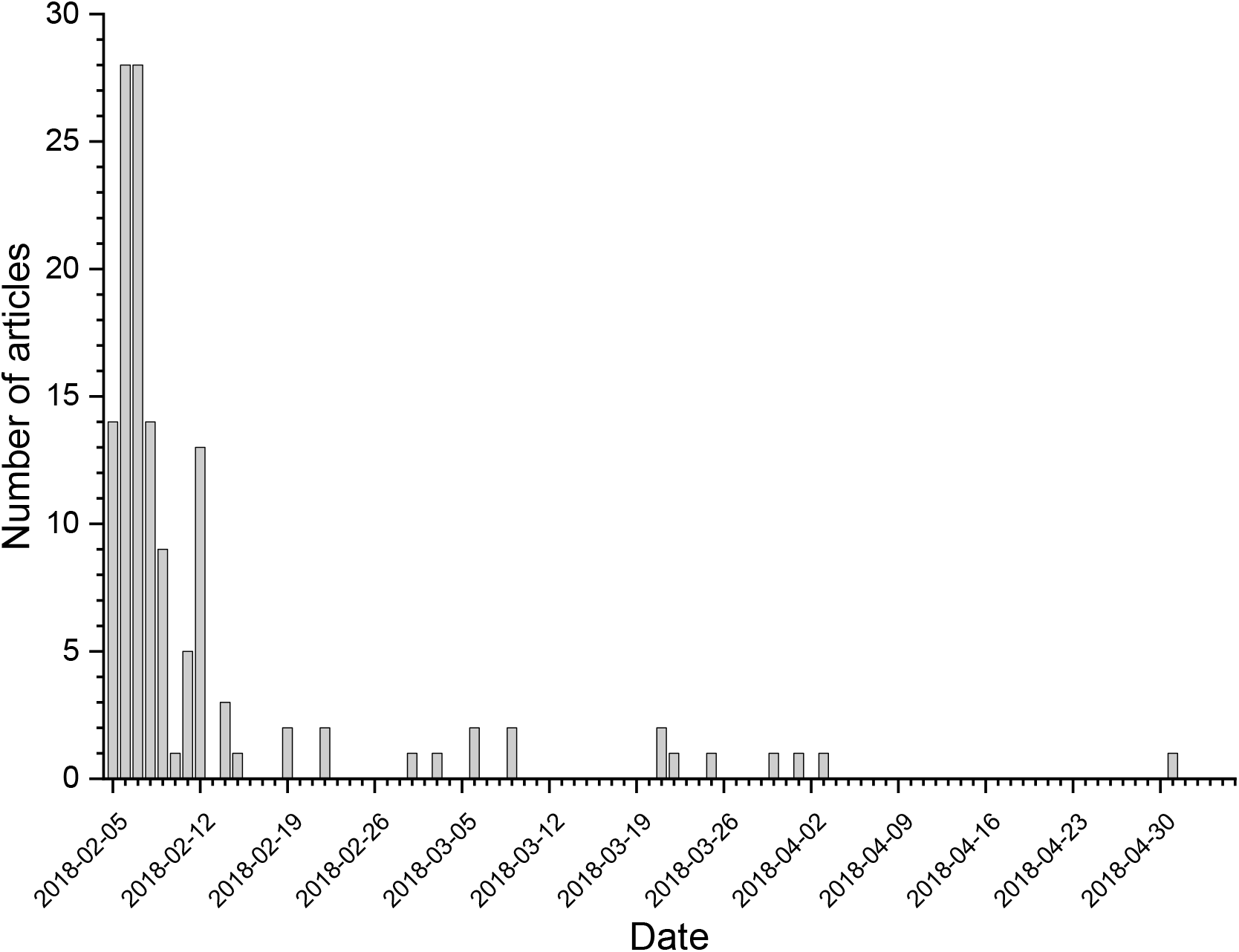
Histogram of publication dates of article covering marbled crayfish paper by Gutekunst et al. (2018).

A *New York Times* article appearing on the same day the paper was released was responsible for a cascade of additional coverage. The *New York Times* article was cited as a source in 25.6% of articles (34 of 133). A comparable story in *The Atlantic*, released on the same day, was cited as a source in 3.8% of articles (5 of 133), three of which also cited the *New York Times*. The *New York Times* article was also reprinted in four other venues, with different headlines in each.

## DISCUSSION

The media coverage following publication of a significant new paper about marbled crayfish (Gutekunst et al. 2018) were framed as “news,” not “explainers” of established, well understood facts that a reader might not happen to know. But the news headlines in this study rarely conveyed new information, instead focusing on facts that was known years ago. The most commonly mentioned point in headlines, that marbled crayfish reproduce by cloning, was known for over a decade (Scholtz et al. 2003; Martin et al. 2007).

There are at least two negative consequences in old research findings being presented as though they are news. First, it underplays the importance of incremental advances in knowledge gained through routine scientific work. The headlines make it appear that the scientific understanding of marbled crayfish emerged at single time, i.e., they convey a “breakthrough” narrative. But the reality is that the knowledge about marbled crayfish represents a decade and a half of research performed by many teams. Furthermore, because scientists read news headlines, this may make scientists more likely to cite Gutekunst et al. (2018) for advances that they did not make, effectively draining citations (and associated academic credit and prestige) from other authors.

On the positive side, the presentation of old information in headlines mean that media coverage of new scientific papers gives older papers a “second chance” (or third, etc.) to reach an audience, and thus provides opportunities for people to become acquainted with basics.

The focal paper (Gutekunst et al. 2018) specifically reported on the invasiveness of marbled crayfish in Madagascar, but more headlines specifically mentioned the presence of marbled crayfish Europe (12.7% of article headlines) than Madagascar (1.5%). While it is not factually wrong to say that marbled crayfish are spreading throughout Europe – they are (Nonnis Marzano et al. 2009; Chucholl and Pfeiffer 2010; Martin et al. 2010b; Chucholl et al. 2012; Bohman et al. 2013; Samardžić et al. 2014; Novitsky and Son 2016; Pârvulescu et al. 2017) – such headlines perpetuate bias. Centering narratives on Europe imply that rich, nations predominantly inhabited by white people are more newsworthy and important than nations that are not.

This study also clearly shows the importance of science reporting in key established media sites. About a quarter of all coverage cited a *New York Times* article, which suggests it triggered the mention rather than the technical paper itself or the press release. This suggests that scientists who are genuinely interested in broad public outreach should specifically target journalists with connections to large established media outlets.

The coverage of this story suggests that science reporting has a weekly “news cycle,” Nine days after the story ran, there were never more than 2 new articles in a day. This is consistent with the weekly publication schedules of “flagship” scientific journals, such as *Nature* and *Science*., which often prompt significant media coverage. Thus, journalists may be used to working in a weekly time frame. But there was a lengthy “tail” of coverage: new, original articles continued to appear almost three months after the initial publication of the research paper.

Accuracy of scientific information is often a main concern for scientists and journalists. But as this study shows, science headlines can be accurate, in the sense that they present correct facts, but still misleading. Old facts are presented in headlines for articles that are supposed to focus on new facts. Facts that are not in a research paper appear in headlines that are ostensibly about it, downplaying the achievements of other authors and distorting the perception of scientific progress.

